# De *novo* protein structure prediction using ultra-fast molecular dynamics simulation

**DOI:** 10.1101/262188

**Authors:** Ngaam J. Cheung, Wookyung Yu

## Abstract

Modern genomics sequencing techniques have provided a massive amount of protein sequences, but experimental endeavor in determining protein structures is largely lagging far behind the vast and unexplored sequences. Apparently, computational biology is playing a more important role in protein structure prediction than ever. Here, we present a system of *de novo* predictor, termed *NiDelta*, building on a deep convolutional neural network and statistical potential enabling molecular dynamics simulation for modeling protein tertiary structure. Combining with evolutionary-based residue-contacts, the presented predictor can predict the tertiary structures of a number of target proteins with remarkable accuracy. The proposed approach is demonstrated by calculations on a set of eighteen large proteins from different fold classes. The results show that the ultra-fast molecular dynamics simulation could dramatically reduce the gap between the sequence and its structure at atom level, and it could also present high efficiency in protein structure determination if sparse experimental data is available.

---

IN modern biology and medicine, it is a major challenge to determine a protein tertiary structure from its primary amino acid sequence, and it has significant and profound consequences, such as understanding protein function, engineering new proteins, designing drugs or for environmental engineering (Röthlisberger *et al*. 2008; Davis and Baker 2009; Qian et *al*. 2004). Nowadays, more and more protein sequences are being produced by genomics sequencing techniques. Despite tremendous efforts of community-wide in structural genomics, protein structures determined by experiments, such as X-ray crystallography or NMR spectroscopy, cannot keep the pace with the explosive growth of protein sequences (Ovchinnikov et *al*. 2017). Since it requires numerous time and relatively expensive efforts, experimental determination of protein structures is lagging behind, and the gap between sequences and structures is widening rather than diminishing (Marks et *al*. 2012).

Amino acid sequences contain enough information for specifying their three-dimensional structures (Anfinsen 1972), thus which provides the principle for predicting three-dimensional structure from sequence. Accordingly, in the past decades, computational prediction of protein structures has been a long-standing challenge, and a number of computational methods have been contributed to bridge the gap, which may be able to be reduced or filled if the approaches can provide predictions of sufficient accuracy (Marks et *al*. 2012). As efficient models, template or homology modeling methods (Sali and Blundell 1993; Kinch et *al*. 2016; Zhang et *al*. 2016) utilize the similarity of the query sequence (target) to at least one protein of known tertiary structure, and protocols in these methods enable to accurately predict protein three-dimensional conformation from its amino acid sequence. However, template or homology models cannot work if there is no determined structure in the same protein family as that of the query sequence. Only relying on the amino acid sequence and no structural template, *de novo* approaches depend on an effective conformation-searching algorithm and good energy functions to build protein tertiary structures.

Nowadays, *de novo* predictors remain restricted to small proteins, and most of them are extremely difficult to achieve on large proteins because of the vast conformational space and computational bottlenecks (Das and Baker 2008; Shen and Bax 2015). Some of these *de novo* approaches rely on assembling proteins from short peptide fragments, which are derived from known proteins based on the sequence similarity (Kinch *et al*. 2016; Zhang *et al*. 2016). For example, Rosetta utilizes sequence-similar fragments by searching against three-dimensional structure databases followed by fragment assembly using empirical intermolecular force fields (Bradley *et al*. 2005). Although many striking *de novo* advances have been achieved, such methods have worked on smaller proteins that have less than 100 amino acids (Kim et *al*. 2009; Söding 2017), unfortunately, the de *novo* structure prediction problem is still unsolved and presents a fundamental computational challenge, even for fragment-based methods (Kim et *al*. 2009).

Here we describe an approach, termed *NiDelta*, to predict protein tertiary structure from amino acid sequence. *NiDelta* models a protein structure from its amino acid sequence primarily involving three steps: (a) predicting torsional angles (*ϕ*, *ψ*) based on the convolutional neural network (CNN); (b) capturing residue contacts based on evolutionary information; and (c) sampling conformation space by ultra-fast Molecular Dynamics simulation.

## 1. Materials and Methods

In this section, the developed *NiDelta* is described in details. The framework of *NiDelta* is illustrated in Fig. 1. As shown, for a given target sequence, *NiDelta* will prepare two main constraints for launching a coarse-grained molecular dynamics (CGMD) — *Upside* (Jumper *et al*. 2016) for sampling conformation space. Firstly, we construct a non-redundant sequence database to building a deep convolutional neural network (LeCun *et al*. 1999) (termed *Phsior*, a module in Sibe web-server (?) for predicting torsional angles (*ϕ*, *ψ*) of a given query amino acid sequence. Thereafter, for the same sequence, we search it against UNIREF100 database (Suzek *et al*. 2015) by HMMER (Eddy 2011) to obtain an alignment of multiple sequences. Accordingly, residue contacts are inferred from the multiple sequence alignment, which encodes co-evolutionary information contributing to coupling relationship between pairwise residues. Then the *Upside* (Jumper *et al*. 2016) is launched for protein conformation samplings with the constraints of predicted torsion angles based on convolutional neural network and contacts derived from evolutionary information.

**Fig. 1.**
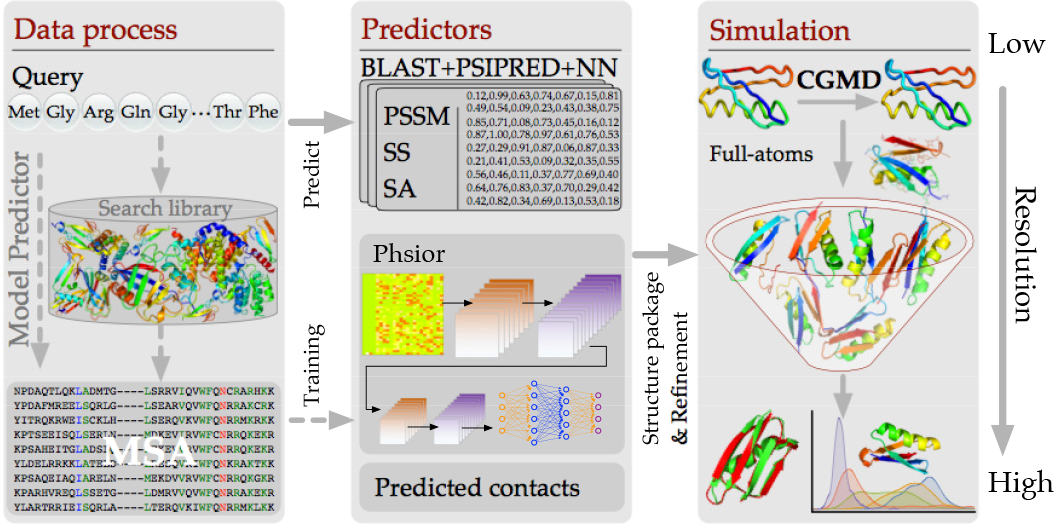
The system flowchart that is used for predicting protein tertiary structure. At the first stage, *NiDelta* constructs both training dataset and MSA for *Phsior* and residue-contacts estimator, respectively. The predicted torsion angles (*ϕ*, *ψ*) and estimated residue-contacts are used as constraints for parallelly launching 500 *Upside* simulations, each of which starts with an extended model represented by a simplified structure for sampling its conformation space.

### A. Torsional angles prediction

The benchmark dataset for *Phsior* is collected from RCSB PDB library and pre-culled through PISCES. The dataset is of 50% sequence identity, 1.8 Å resolution, and 0.25 R-factor (November 6, 2017). In the dataset, there are 10,586 chains used as the sequence library. The experimental values of the (*ϕ*, *ψ*) angles are extracted by STRIDE program (Frishman and Argos 1995), and the N- and C-terminal residues are neglected because of the incompleteness of four continuous backbone atoms (Wu and Zhang 2008).

*Phsior* is a real-value predictor developed based on the convolutional neural network for predicting the torsion angles (*ϕ*, *ψ*). Briefly, the architecture of *Phsior* is illustrated in Fig. 2 (see also Supplementary Methods). *Phsior* extracts three types of sequence-based features involving position-specific scoring matrices (PSSM), secondary structure (SS), and solvent accessibility (SA). The PSSM is generated by PSIBLAST (Altschul *et al*. 1997) search of the query against anon-redundant sequence database with 20 log-odds scores taken at each position. The secondary structure (SS) is predicted by PSI-PRED (Jones 1999), with the three states defined as alpha-helix, beta-strand, and coil. The solvent accessibility (SA) is predicted by the neural networks (Chen and Zhou 2005). These three kinds of features will be normalized and used as inputs of the CNN model.

**Fig. 2.**
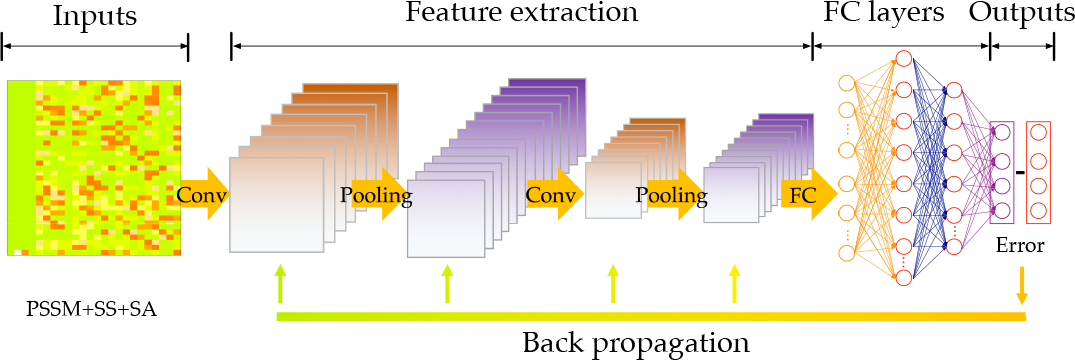
The architecture of *Phsior*. The feature extraction stage includes convolutional and max-pooling layers. The first convolutional layer consists of several 5-filters, which slide along the input feature matrix. The second convolutional layer works on successive convolutions from previous layers. Following the filters, two fully connected layers are presented to integrate and make final predictions of *ϕ* and *ψ*.

*Phsior* begins with a simplistic baseline to predict torsion angles (*ϕ*, *ψ*) by employing a fixed-size context window of 17 amino acids through two convolutional layers and two fully-connected layers (as illustrated in Fig. 2). *Phsior* predicts the torsion angles (*ϕ*, *ψ*) of the central amino acid via the final fully-connected layer.

As inputs of the deep network, data is normalized to the range of 0.0 to 1.0. Then we use a window size of 17 to include the neighborhood effect of close amino acids. The data produces a probability map of 35 × 24. The convolutional layers in *Phsior* are to detect recurrent spatial patterns that best represent the local features, while max-pooling layers are to down-sample the features for increasing translational invariance of the network. The fully connected layers are to integrate for the outputs and then make the final predictions for the torsion angles (*ϕ*, *ψ*
).

In *Phsior*, a convolutional filter can be interpreted as sliding along the input feature matrix, sharing and/or re-using the same few weights on each local patch of the inputs. Fig. 2 illustrates the convolutional layers working on an example amino acid from training samples. In particular, the first convolutional layer in Fig. 2 consists of a 5-filter which is repeated several times as it slides along the feature matrix. Generally, local properties of the input data are important, the small filters show their capability in learning and maintaining information derived from the amino acid sequence at different scales.

In the output layer of *Phsior*, sine and cosine are employed to remove the effect of angle periodicity. Predicted sine and cosine values are converted back to angles by using the equation *α* = tan^-1^ [sin (*α*) / cos(*α*)].

Weights of *Phsior* are randomly initialized according to a zero-centered Gaussian distribution with a standard deviation of 5/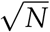 (N is the number of inputs in each layer).

### B. Residue contact prediction

Recently, residue-contacts lead *de novo* prediction in a fast progress, like direct coupling analysis (DCA) (Marks *et al*. 2011; Morcos *et al*. 2011), protein sparse inverse covariance (PSICOV) (Jones *et al*. 2011) or Gremlin (Balakrishnan *et al*. 2011; Kamisetty *et al*. 2013) those are all able to disentangle such indirect correlations, and extract direct coevolutionary couplings. These have been found to accurately predict residue-residue contacts — provided a sufficiently large MSA.

Co-evolutionary information encoded in the amino acid sequences highly contributes to residue contacts (Marks *et al*. 2011; Morcos *et al*. 2011; Jones *et al*. 2011; Balakrishnan *et al*. 2011; Kamisetty *et al*. 2013). Accordingly, we estimate pairwise residue contacts from protein multiple sequence alignment (MSA). Firstly, we prepared the MSAs for each studied protein by searching the query sequence against the UniRef100 database (Suzek *et al*. 2015) using the jackhmmer method (Eddy 2011). The obtained MSAs were trimmed based on a minimum coverage, which satisfies two basic rules: (1) a single site with more than 50% gaps across the MSA will be removed; and (2) the percentage of aligned residues between the query and the obtained sequence less than a given threshold will be deleted from the MSA.

After filtering the MSA, we start to estimate coupling scores between pairwise residues according to the direct coupling analysis (DCA) algorithm (Weigt *et al*. 2009; Morcos *et al*. 2011; Marks *et al*. 2011, 2012). Given the MSA, we can easily compute the single site frequency *f_i_* (*A_i_*) and joint frequency *f_i__j_*(*A_i_, A_j_*). To maximize the entropy of the observed probabilities, we can calculate the effective pair couplings and single site bias to meet the maximal agreement between the distribution of expected frequencies and the probability model of actually observed frequencies.

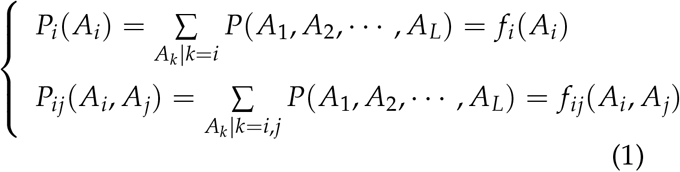

Maximizing the entropy of the probability model, we can get the statistical model as follows,

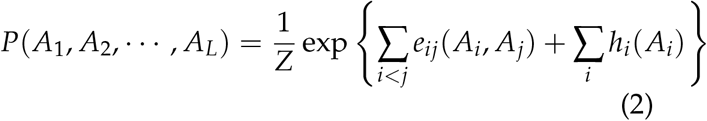

where *Z* is a normalization constant, e_ij_ (·, ·) is a pairwise coupling, and *h_i_*(·) is a single site bias.

Accordingly, the mathematical definition of the score in DCA approach is formulated as follows,

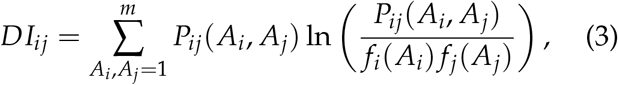

where *DI_ij_* is the direct coupling score between pairwise amino acids at the ith and jth sites in the MSA. The top-ranked set of *DI_ij_* are converted to contacts between pairwise residues (Marks *et al*. 2011).

### C. Ultra-fast molecular dynamics simulation

In the proposed method, we launched a coarsegrained molecular dynamics simulation (CGMD, termed *Upside)* (Jumper *et al*. 2016) for sampling the conformation space of a given target sequence. In the *Upside*, the model is presented by a reduced chain representation consisting of the backbone N, C*_α_*, and C atoms. The *Upside* launches dynamics simulations of the backbone trace including sufficient structural details (such as side chain structures and free energies). The inclusion of the side chain free energy highly contributes to the smooth the potential governing the dynamics of the backbone trace (Jumper *et al*. 2016).

In this study, the predicted torsion angles (*ϕ*, *ψ*) and the inferred residue contacts are used as constraints to run *Upside* simulations from an extended structure. In the Upside, the Miz-Jern potential is employed without using the multi-position side chains (refer to (Jumper *et al*. 2016) for more details). For the ith residue, we provide a range for both *ϕ_i_* and *ψ_i_*, such as *ϕ_i_ ∈ [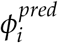* — 20°, 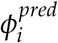 + 20°] and *ψ_i_ ∈[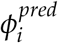 —* 20°, 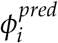 + 20°]. This strategy guides the *Upside* sample the Ramachandran map distribution for the secondary structures. On the other hand, the contacts provide distant constraints for pairwise residues in spacial, which contribute to sample the tertiary structures. According to the design of experiment conducted, we select top 2L residue contacts (distant of C*β*-C*β* between pairwise residues less than 7.5Å) as constraints. The *Upside* is configured by setting weights for hydrogen-bond energy, side chain radial scale energy, side chain radial scale inverse radius and side chain radial scale inverse energy to −4.0, 0.2, 0.65 and 3.0, respectively. For, each protein sequence, we launched 500 individual simulations starting from the same extended conformation with a duration time of 500,000 and capture conformations at every 500 frames.

## 2. Results and Discussion

As described in the methods, we sought to provide a template-free prediction system for folding proteins. The approach only depends on sequence information without any structural templates or fragment libraries. We demonstrate the predictive ability of the developed system on a set of candidate structures of proteins over a range of protein size and different folds. The details of eighteen proteins are reported in Table 1. As illustrated in the table, we present the protein name, PDB id in RCSB database, length of each protein sequence, protein folds, the number of sequences in each MSA, centroid and best C*_α_*-RMSD with corresponding TM-score (computed by TMscore software (Zhang and Skolnick 2004)). All the comparisons of C*_α_*-RMSD and TM-score are computed in full length of each target protein.

**Table 1.** Accuracy of predicted proteins.

| Protein name | L<sup>a</sup> | Fold | N<sup>b</sup> | C<sub>α</sub>-RMSD<sup>c</sup><sub>crt</sub> | C<sub>α</sub>-RMSD<sup>d</sup><sub>best</sub> | Ref. PDB |
| --- | --- | --- | --- | --- | --- | --- |
| CrR115 | 134 | $\alpha/\beta$ | 6.0k | 4.57 (0.60) | 2.51 (0.79) | 2lcgA |
| ER553 | 141 | $\alpha/\beta$ | 98k | 4.11 (0.67) | 3.11 (0.76) | 2k1sA |
| C-H-RAS P21 | 166 | $\alpha/\beta$ | 574k | 4.08 (0.75) | 2.98 (0.77) | 5p21A |
| HR2876B | 107 | $\alpha/\beta$ | 6.9k | 4.52 (0.64) | 3.42 (0.69) | 2ltmA |
| CG2496 | 115 | $\alpha/\beta$ | 19.8k | 2.80 (0.75) | 2.19 (0.80) | 2kptA |
| Thioredoxin | 105 | $\alpha/\beta$ | 214k | 2.88 (0.73) | 2.12 (0.80) | 1rqmA |
| CheY | 130 | $\alpha/\beta$ | 887k | 8.08 (0.57) | 4.21 (0.64) | 1e6kA |
| Ribonuclease HI | 143 | $\alpha/\beta$ | 63.8k | 9.46 (0.42) | 5.47 (0.56) | 1f21A |
| Isomerase | 108 | $\alpha + \beta$ | 68.4k | 5.17 (0.57) | 3.34 (0.68) | 1r9hA |
| OR36 | 134 | $\alpha/\beta$ | 6.2k | 6.42 (0.47) | 4.08 (0.68) | 2lciA |
| MTH1958 | 136 | $\beta$ | 43.9k | 7.94 (0.37) | 4.77 (0.63) | 1tvgA |
| SgR145 | 173 | $\alpha/\beta$ | 771k | 6.87 (0.51) | 4.99 (0.63) | 3merA |
| Tpx | 167 | $\alpha/\beta$ | 185k | 3.03 (0.77) | 2.38 (0.83) | 2jszA |
| YwIE | 150 | $\alpha/\beta$ | 40.6k | 3.42 (0.76) | 2.52 (0.82) | 1zggA |
| FluA | 173 | $\beta/\alpha$ | 15.9k | 7.09 (0.50) | 5.02 (0.59) | 1n0sA |
| Rhodopsin II | 222 | $\alpha$ | 3.4k | 5.68 (0.64) | 5.24 (0.65) | 2ksyA |
| Savinase | 269 | $\alpha/\beta$ | 102k | 6.83 (0.65) | 5.17 (0.69) | 1svnA |
| MBP | 370 | $\alpha/\beta$ | 200k | 8.85 (0.51) | 6.49 (0.64) | 1dmbA |

<sup>a</sup> Protein length.<sup>b</sup> Number of sequences obtained by jackhmmer method.<sup>c</sup> RMSD in full length of the centroid structure of the largest cluster compared to the native shown in Å(TM-score).<sup>d</sup> RMSD in full length of the best structure compared to the native shown in Å (TM-score).

We first compare the predictions on the torsion angles (*ϕ*, *ψ*) of the target proteins listed in Table 1 among Anglor (Wu and Zhang 2008), Spider2 (Hef-fernan *et al*. 2015), and our model *Phsior* over the eighteen target proteins. For a fair comparison, a criterion is defined by the mean absolute error (MAE) to validate the predicted angles (*ϕ*, *ψ*), and the MAE is to measure the average absolute difference between the experimentally determined and predicted angles.

Accordingly, the MAE is formulated as follows,

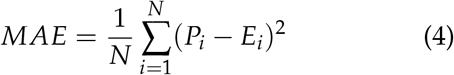

where *N* is the number of residues (excluding N- and C-terminals) in a protein. *P_i_* is the predicted value for *i*th residue, and *E_i_* is the experimental value of *j*th residue in the protein.

As illustrated in Fig. 3 (see also Fig. S1), the proposed *Phsior* and Spider2 (Heffernan *et al*. 2015) are in comparable performances on the target proteins listed in Table 1. They were all better than those of Anglor (Wu and Zhang 2008). The MAE of torsion angle(*ϕ*, *ψ*) predicted by Anglor on each protein was almost three times of that of *Phsior* and Spider2, especially on the transmembrane protein Rhodopsin II (PDB ID: 2KSY), the difference remains the largest among all the comparisons. As we know, Anglor is a combined predictor of support vector machine and simple feedforward artificial neural network, while *Phsior* and Spider2 are based on the deep neural network. Accordingly, the better performances could be a result of the powerful capability of the deep learning technique. Although *Phsior* was slightly better than that of Spider2 on several benchmark targets, it seems that *Phsior* is more stable on the predictions.

**Fig. 3.**
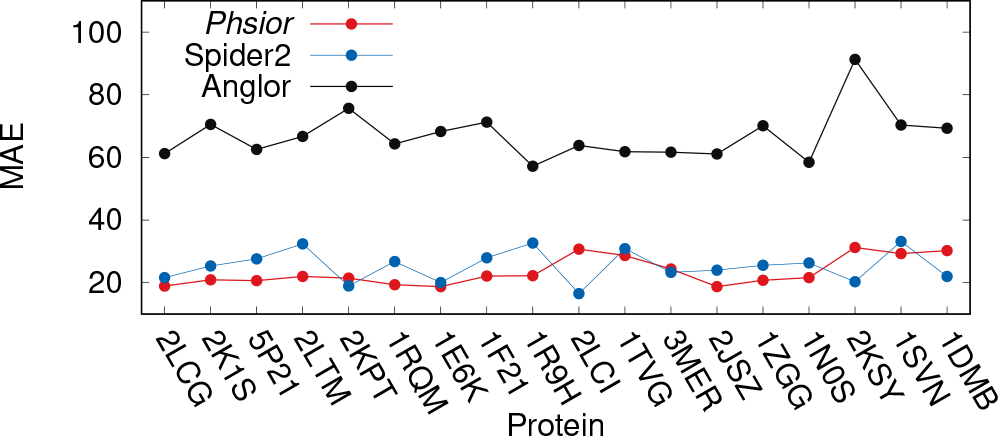
Comparison on the MAE of the predicted torsion angles (*ϕ*, *ψ*) among Anglor, Spider2, and *Phsior*.

Since the residues in a region of protein chain are more likely to be related than independent amino acid far away, this ‘locality’ make the prediction ability of the CNN method more powerful. The CNN model can capture the dependences of amino acids in the same chain, which can result in much information of ‘locality’ among resides. Moreover, the proposed strategy of the predicted torsion angles (*ϕ*, *ψ*) can guide the *Upside* to efficiently sample conformation space at high speed. Accordingly, in the developed system, the predictions of *Phsior* are preferred and used as constraints in the *Upside*.

The quality of the predictions by *Phsior* is roughly good to contribute to the constraints for the *Upside* simulation, although there were also several not so good predictions (worse than those of Spider2). However, this did not mean that we could not simply to use the predicted torsion angles (*ϕ*, *ψ*) as starting for the *Upside* simulation. Instead, we found it efficient to pre-defined a range for each torsion angle (Supplementary Methods).

We further investigate whether co-evolving sequences can provide sufficient information to specify a good model for assessing blind predictions of protein tertiary structures close to the crystal structures. The predicted residue-contacts mostly correlated with the native ones. However, the inferences from the MSA always included noises and false positive predictions, which meant that they could not be simply used for the *Upside*. Instead, we found it efficient and important to generate a potential by sigmoid-like function for the *Upside* (see also Supplementary Methods).

For the most of 18 proteins, the estimated residue-contacts include several sparse but informative true positive predictions, making them useful constraints for the *Upside* sampling. Only for the protein OR36 (PDB ID: 2LCI) did NiDelta fail to infer a residue-contact map (Fig. S2), this could result from less diversity in its MAS. Although the bad residue-contacts occur, the *Upside* can robust to the noises to perform simulation based on Ramachandran map distribution. This could result from the strategy designed for the predicted torsion angles (*ϕ*, *ψ*).

As shown in Fig. 4, nine representative residue contacts estimated from the MSAs present to compare to the corresponding native ones (see also Fig. S2). The estimated residue-contacts include noises, which (significantly incorrect predictions) are highlighted in green circles in Fig. 4. For instance, there are five groups of incorrect predictions (noises) in the inferred residue-contacts of the HR2876B protein (PDB ID: 2LTM). The noises possibly led the mis-folding of the unstructured regions of the protein as shown in Fig. 6. The similarity can also be found in the Thioredoxin (PDB ID: 1RQM) and the YwIE (PDB ID: 1ZGG) proteins.

**Fig. 4.**
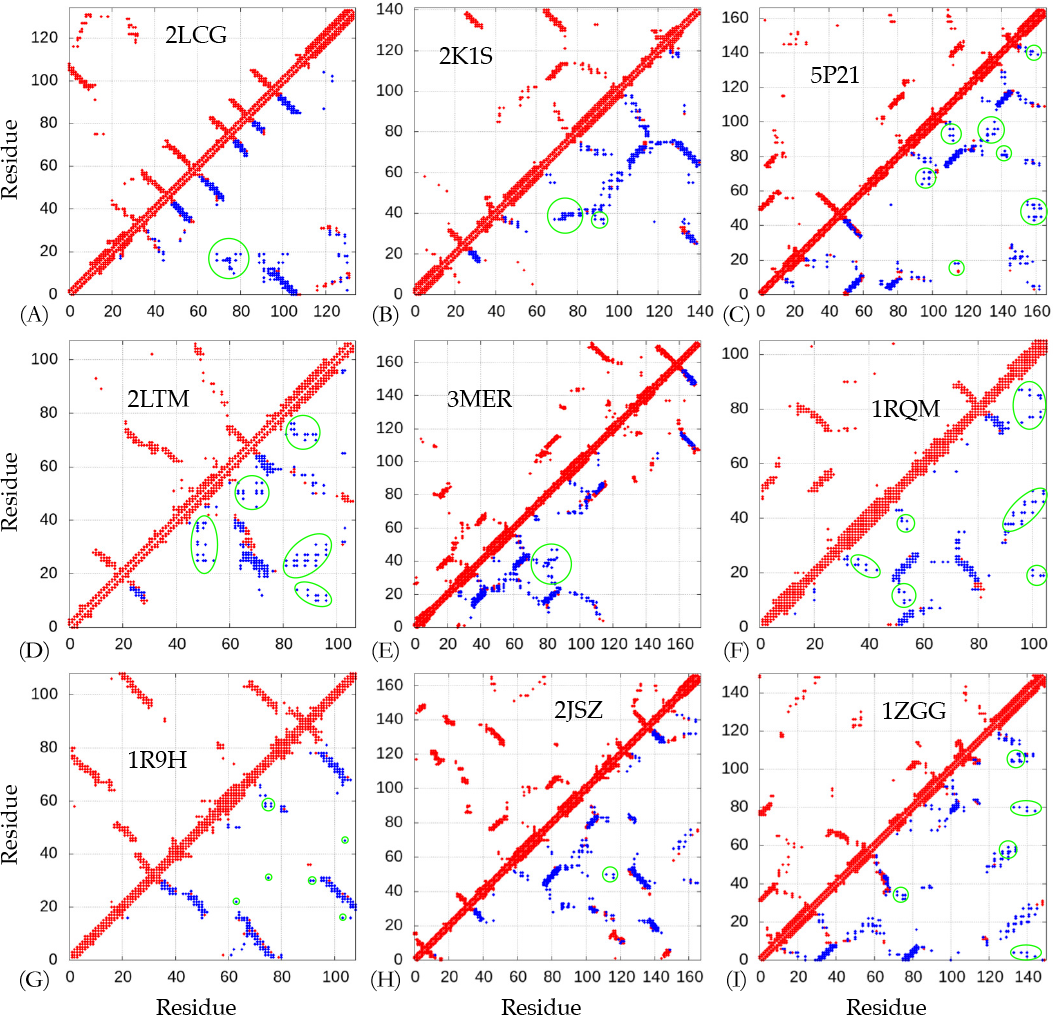
The predicted residue-contacts for highlighted targets. All the residue-contacts (top 2L) used in the *Upside* simulations are shown in blue filled squares. The native and estimated residue-contacts are in red and blue, respectively. The dots in green circles are noises (false positive inferences).

Immediately after predicting the torsion angles and residue-contacts, it is usually straightforward to assign the ranges for the angles (*ϕ*, *ψ*) and the potentials for interactions between pairwise residues, respectively. Then we launch the ultra-fast coarsegrained molecular dynamics *(Upside* (Jumper *et al*. 2016)) with the restraints of predicted torsional angles and residue contacts (Supplementary Methods).

For each protein sequence, 500 *Upside* simulations (trajectories) were performed, starting from the unfolded structure. We collected the trajectories for analyzing, and last 50 structures captured from each simulation trajectory were selected from 500 trajectories for clustering (total number is 25,000). We conducted a clustering analysis of the structures using *fast_protein_cluster* software (Hung and Samudrala 2014) to cluster the structures and calculate the tightness of those clusters, which represent conformational ensembles predicted from each protein sequence. For further study, centroids of the top 5 clusters were selected as our "blind predicted models". The clustering results are illustrated in Fig. 5. The biggest cluster has the strongest tightness on the most target proteins (except proteins CG2496, CheY, Ribonuclease HI and Savinase).

**Fig. 5.**
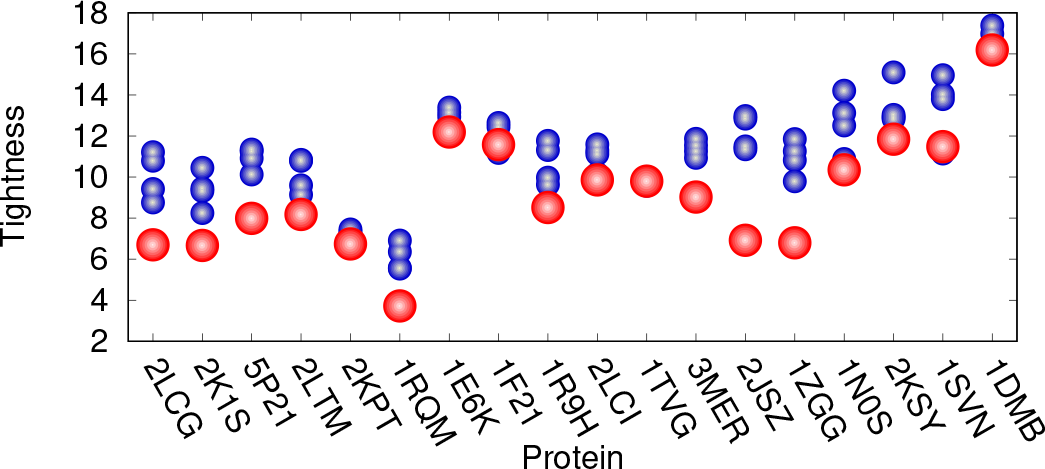
Top five clusters of each target proteins listed in Table 1. The biggest clusters are colored in red, while other clusters are represented in blue.

To visualize how the structural agreement between the predicted models and the native structure, for nine representative cases, we plotted the proteins corresponding to the best predictions against their C*_α_*-RMSD relative to the experimental reference structures (Fig. 6, and see also Fig. S3). As illustrated in Fig. 6, structural results of the *NiDelta* for nine representative test proteins. In the figure, ribbon models of the lowest C*_α_* — *RMSD* structure (green) (calculated with the *Upside*) superimposed on the corresponding experimental structure (red). For example, as an interesting representative, the C-H-RAS P21 protein p21 (PDB ID: 5P21) involves in a growth promoting signal transduction process (Barbacid 1987). As shown Fig. 4(C), al were noisy predictions in the restraints of torsion angles (*ϕ*, *ψ*) (Fig. 3 and Fig. S1) and residue-residue contacts (Fig. 4(C)), The best C*_α_*-RMSD of 3 Å model of the C-H-RAS P21 protein is in the same fold with TMscore of 0.76, and also the centroid model of the largest cluster is blind prediction of C*_α_*-RMSD of 4.1 Å and TMscore of 0.75, which indicates that the Upside can is able to fold a large protein and robust to the noises although the existing noises may mislead the simulation in sampling its tertiary structure (e.g. the prediction of the OR36 protein, see Fig. S2(I) and S3). As illustrated in Fig. 6(F), the structure of the Thioredoxin protein (PDB ID: 1RQM) consists of a central core of a five-stranded *β*-sheet surrounded by four exposed *α*-helices (Leone *et al*. 2004). Although the noises and false positive predictions exist in residue contacts (Fig. 4), the predicted model of the best C*_α_*-RMSD is 2.1Å, and its corresponding TMscore is as high as 0.8, which mean that the model is almost structurally identity to the native fold. The successful predictions can be also found in the centroid model in top 1 cluster of the C*_α_*-RMSD is 2.9Å and TM-sore 0.73 (Table 1). The blind predictions obtained from the clustering results show that most of the 500 folding simulations converged to similar groups with strength tightness (Fig. 5). This could result from that the *Phsior* providing more accurate angles (*ϕ*, *ψ*) help the *Upside* robust to the noises and inaccurate information. As shown in Fig. 6(I) (red), the tertiary fold of the YwlE protein (PDB ID: 1ZGG) is a twisted central four-stranded parallel *β*-sheet with seven *α*-helices packing on both sides, in which the active site is favorable for phosphotyrosine binding (Xu *et al*. 2006). The results of the YwlE protein in Fig. 4(I), Fig. 3, and Fig. 5 further demonstrate that *Upside* has a strong predictive ability in folding a protein with inaccurate constraints, even with incorrect information.

**Fig. 6.**
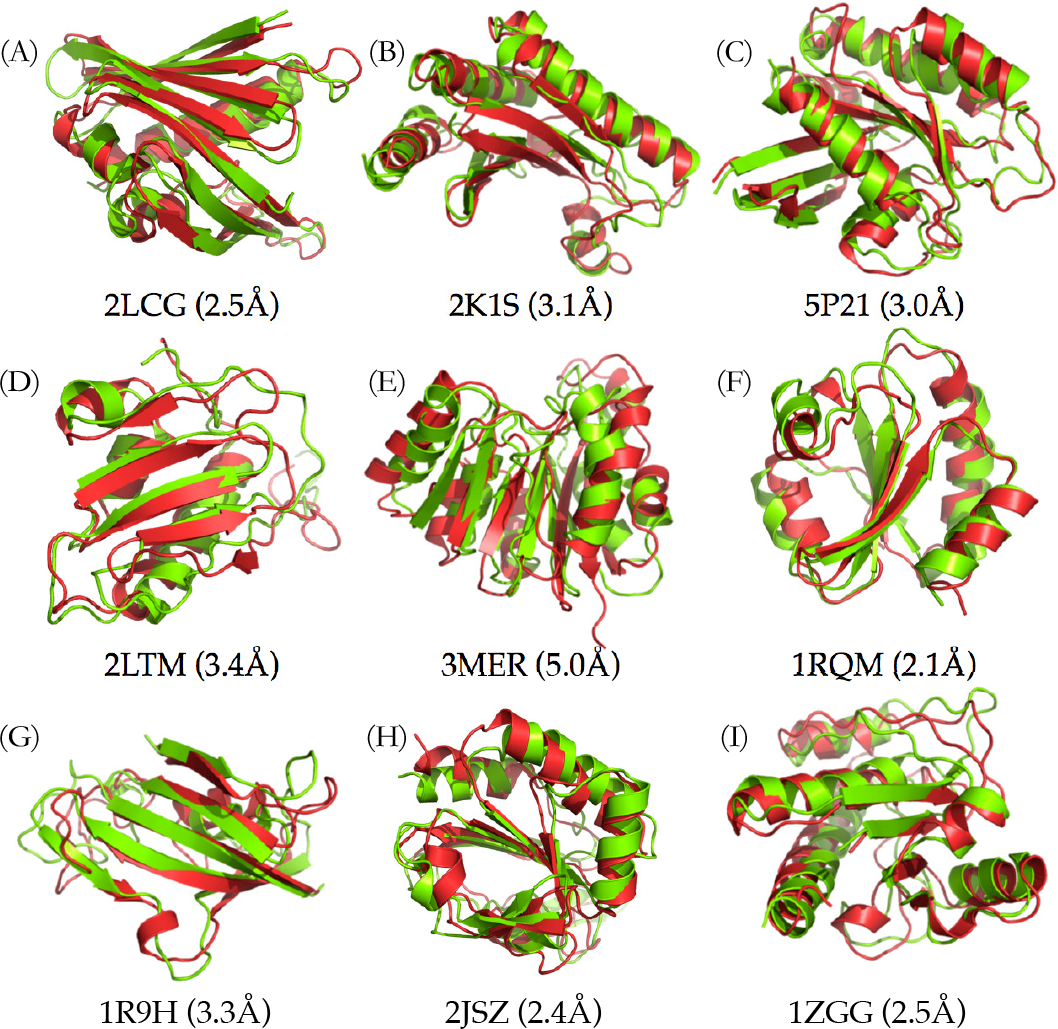
Highlighted predicted structures. Visual comparisons on nine of the target proteins (the native and predicted structures are in red and green, respectively).

Three models (three proteins of more than 200 residues) corresponding to each of the centroid of the biggest clusters are illustrated in Fig. 7. The C*_α_*-RMSD values of the centroids compared to the known structures are 5.7Å, 6.8Å, and 8.9Å for Rhodopsin II, Savinase, and MBP proteins, respectively. The protein Rhodopsin II is a membrane protein predicted by the proposed system. For the top ranked predicted model (5.2Å C*_α_*-RMSD with full length alignment, as shown in the center in Fig. 7(A)), the terminal helix is misaligned, but the orientations of other six helices are in an excellent agreement with those of the crystal structure. As illustrated in the right of Fig. 7(B), the centroid model is also misaligned in the terminal helix, but it provided more structural details as shown in the helices 5 and 6. The structure of the Savinase protein chosen as the protein of interests has an *α/β* fold consisting of 9 helices and 9 strands, which is a representative of subtilisin enzymes with maximum stability and high activity (Betzel *et al*. 1992). The model of the best C*_α_*-RMSD has correct topography of seven *β*-strands and eight *α*-helices, while there are six *β*-strands and seven *α*-helices in the centroid model. Flexibility in the conformation occurs in the C-terminal region of Savinase protein (Betzel et *al*. 1992), which make the prediction particularly challenging. As shown, both the models of the best C*_α_*-RMSD and centroid capture the structural information. As shown in Fig. 7(C), the largest protein tested in the benchmark test is the maltodextrin binding protein (MBP), which is from Escherichia coli serving as the initial receptor for both the active transport of and chemotaxis toward a range of linear maltose sugars (Sharff *et al*. 1993), with 370 amino acids. It is significantly larger than proteins that can be predicted by other de *novo* computational approaches (Marks et *al*. 2011). With the predicted angles (*ϕ*, *ψ*) and residue-contacts, the *Upside* can achieve a blind model of C*_α_*-RMSD 8.9Å and TM-score 0.51, which indicates that the model is in about the same fold (Zhang and Skolnick 2004) and efficiently predictive ability of the proposed approach in the particularly challenging *de novo* structure prediction of large proteins. Accordingly, a strength of the proposed method is demonstrated here is that, based on the centroids of those top 5 clusters, we can potentially develop iterative predictions for larger proteins by collecting centroid models and extracting the informative constraints from previous round of simulations as refinements.

**Fig. 7.**
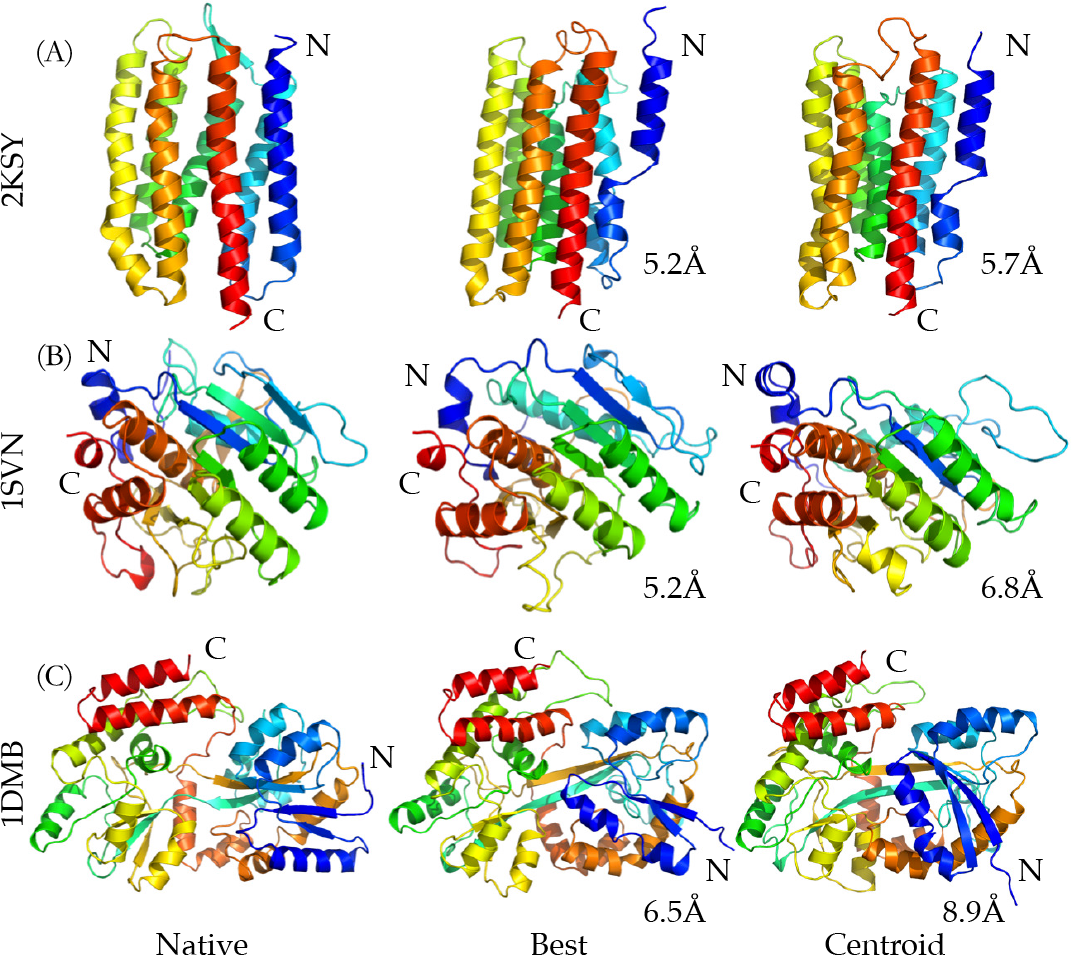
Visual comparisons on three target proteins with more than 200 residues. The highlighted structures from left to right are the native, the structures of the best C*_α_*-RMSD, and the centroid of the biggest cluster, respectively.

## 3. Conclusion

This study presents a way of integrating predicted torsion angles & residue contacts within an ultra-fast molecular dynamics simulation (*Upside*) to achieve *de novo* structure prediction on large proteins. We have tested the proposed approach on the proteins of more than 100 residues and different folds, and also have achieved the agreement of the predictions with the native structures of the benchmark proteins. Statistically determined residue-contacts from the MSAs and torsion angles (*ϕ*, *ψ*) predicted by deep learning method provide valuable structural constraints for the ultra-fast MD simulation *(Upside)*. The *Upside* provides a simulation with high computational efficiency, which allows users predict structures of large proteins in several CPU hours, get highly accurate models, and details of partial protein folding pathways. Depending on a portion of structural constraints predicted and estimated from the amino acid sequence, the proposed methodology makes the *Upside* a perfect computational platform for de *novo* structure prediction of large proteins.

Although pairwise couplings statistically inferred from protein multiple sequence alignment is a breakthrough in contribution to computational protein structure prediction, there are a number of limitations. For example, residue-residue contacts cannot be estimated if there are no enough as diverse as possible multiple sequences in an align of a protein family. Additionally, even when we have sufficient sequences, the pairwise contacts contain false positive predictions that may result in incorrectly building the 3D structure of a protein. Another limitation, applicable to all existing approaches, is predicting the torsion angles (*ϕ*, *ψ*). It is challenging to accurately predict torsion angles. *Phsior*, designed based on deep convolutional neural network, is able to predict the angles, but it is difficult to make accurate prediction of each pair (*ϕ*, *ψ*). Although we have provided a strategy to handle the inaccurately predicted torsion angles and noised residue-residue contacts, work that of more deep network and iteratively passes information (e.g. averaged torsion angles and contact maps from top 2 structural clusters) collected from previous round of predictions to the next round is currently underway for better predictions of large proteins.

The predicted models (of the best C*_α_*-RMSD and centroid) are consistent with the crystal structures of their natives, and the validation of our approach on eighteen large proteins suggests that the developed approach is capable in efficiently folding large protein based on predicted constraints. Accordingly, we are confident that future refinement of the approach will be successfully applied to very large proteins and complexes when experimental constraints are available, such as chemical shift, sparse nuclear overhauser effect (NOE) and cryo-electron microscopy (cryo-EM) maps. In summary, we introduce a method *NiDelta* as a *de novo* prediction system for large proteins. We hope this approach will find its place in the fields of both the protein structure prediction and determination in the future.

## Acknowledgments

We thank Drs. T.R. Sosnick, K.F. Freed, J.M. Jumper, and S. Wang for help and advice. This work was supported by DGIST start-up fund (No. 2017010068 and No. 2018010089). We also gratefully acknowledge the DGIST Supercomputing and Big Data Center for dedicated allocation of supercomputing time.

